# Dogs as reservoirs of *Escherichia coli* strains causing urinary tract infection in their owners

**DOI:** 10.1101/302885

**Authors:** Peter Damborg, Heidi Gumpert, Laura Johansson, Bimal Jana, Niels Frimodt-Møller, Luca Guardabassi

## Abstract

It is known that humans and pets living together can share the same *Escherichia coli* strain. In this study we assessed the role played by household pets as reservoirs of *E. coli* strains causing urinary tract infection (UTI) in their owners. Fecal swabs from 15 dogs and six cats living with 19 patients with community-acquired *E. coli* UTI were screened by antimicrobial selective plating to detect *E. coli* displaying the same susceptibility profile of the UTI-causing strain. Pet/patient pairs sharing strains with indistinguishable susceptibility and pulsed-field gel electrophoresis (PFGE) profiles were quantitatively screened for fecal carriage of the UTI-causing strain approximately 10 months later using bacterial counts on selective agar supplemented with the relevant antibiotics. Isolates from both time points were characterized by whole-genome single nucleotide polymorphism (SNP) analysis. PFGE revealed indistinguishable *E. coli* within two (11%) pet/patient pairs. In pair A, the UTI-causing strain was detected 10 months later in both the patient (10^8^ CFU/g) and her dog (10^4^ CFU/g). In pair B, only the dog was colonized with the UTI-causing strain upon re-sampling (10^5^ CFU/g), indicating dog-to-man transmission. For both pairs, less than 70 SNPs distinguished any isolate from the first and second sampling. The study shows regular co-carriership of UTI-causing *E. coli* strains between humans and their pets, and indicates that dogs can be a source of human infection. Although final evidence for transmission is lacking, hygiene precautions should be considered by people fraternizing pets. This may be particularly relevant for persons with a compromised immune system.

## INTRODUCTION

Urinary tract infection (UTI) is a common disease in humans, and approximately 80% of acute, uncomplicated community-acquired UTI incidents are caused by *Escherichia coli* (1). Patients are most often self-infected meaning they are infected with strains colonizing their intestinal tract (2). Most strains causing UTI are extraintestinal pathogenic *E. coli* (ExPEC) (3), which is a pathotype characterized by presence of specific virulence genes such as adhesins, toxins, and polysaccharide coatings enhancing pathogenicity outside the intestinal tract (4). Apart from in humans, *E. coli* strains resembling ExPEC based on their genotype have been identified in different animal species, especially dogs (5–7). Various studies have shown that healthy pets (mainly dogs) and humans living together frequently share intestinal *E. coli* strains (8, 9). Furthermore, cases where the family dog or cat was colonized by a strain causing UTI in a human household contact have been reported in the scientific literature, including a case of UTI caused by an extended-spectrum beta-lactamase (ESBL)-producing strain (9–12). However, pet shedding of *E. coli* strains causing UTI in their owners has not been investigated systematically or longitudinally prior to this study.

The aim of this study was to assess the role played by household pets as reservoirs of *E. coli* strains causing UTI in their owners. Dogs and cats owned by UTI patients were screened for the occurrence of *E. coli* having the pheno- and genotypic characteristics of the strain causing UTI in their owner. When a pet was found to carry a strain indistinguishable from that causing infection in the owner, quantitative shedding of the UTI-causing strain by the pet and the owner was followed up after 10 months, and whole-genome sequencing of multiple pet and human isolates was used to infer genomic micro-evolution of the strain shared by the two hosts.

## MATERIALS AND METHODS

### Patient recruitment and sampling of pets

The Department of Clinical Microbiology (DCM) at Hvidovre Hospital, Copenhagen, processes diagnostic specimens from several hospitals and primary care practices in the southern part of Greater Copenhagen and surrounding areas. In the period from February to May 2014, patients diagnosed at DCM with community-acquired UTI caused by *E. coli* were identified on a daily basis via the laboratory management system. Patients infected with putative ESBL-producing *E. coli* (based on resistance to third generation cephalosporins, 3GC) were enrolled in the study (see details below). For each patient infected with a 3GC-resistant strain, we recruited one or two randomly chosen patients infected with *E. coli* susceptible to 3GC. All patients below 85 years were considered eligible; patients above the age of 85 were for the majority living in nursing homes, hence a healthcare rather than a community setting.

Upon consent from a patient’s general practitioner, the patient was contacted by telephone. For patients below 18 years of age, a parent was contacted instead. Patients/parents were asked if they had a dog or cat at home. In case of positive response, they were informed about the study and invited to participate. Patients/parents agreeing to participate were provided with sampling material and a written instruction on the sampling procedure. Dog owners were instructed to dip a cotton swab into a fresh fecal sample while walking their dog. Cat owners were instructed to dip the cotton swab into fresh feces (< 8h old) in litter trays. On the day of collection, swabs were shipped in commercial transport medium (BBL Cultureswab, Becton Dickinson, USA) to the research laboratory at the University of Copenhagen. All samples were collected within two weeks after diagnosis of the UTI in the owner and processed in the laboratory within 48h of collection. The protocol for patient recruitment was approved by the National Committee on Health Research Ethics (journal record: H-4-2013-FSP-071).

### Bacterial isolation and identification

On the day of receipt, fecal swabs from pets were processed using a selective approach in order to enhance detection of potential low-abundant *E. coli* clones causing UTI in the patients. For pets of patients infected with a 3GC-resistant strain, samples were enriched overnight in MacConkey broth with 1 μg/ml cefotaxime at 37°C. Twenty 20 μl of the enrichment was subsequently plated on MacConkey agar containing 1 μg/ml cefotaxime. If growth was observed the next day, one colony representing each morphological type was sub-cultured and stored at −80°C. For pets of patients infected with a 3GC-susceptible strain, a previously described selective direct plating method was used to detect *E. coli* displaying the same resistance profile of the UTI-causing strain (13). In brief, each fecal swab was uniformly streaked on MacConkey agar, and antimicrobial disc(s) (Oxoid, Basingstoke, UK), representing the antibiotics to which the patient isolate was resistant, were applied onto the agar surface. Upon overnight incubation at 37°C, colonies growing in proximity of the discs or within the inhibition zones and displaying different colony appearance were sub-cultured followed by storage at −80°C. All isolates were identified by matrix-assisted laser desorption ionization-time of flight mass spectrometry (Maldi-TOF MS) (Vitek MS RUO; bioMérieux, Marcy-l’Étoile, France) using *E. coli* ATCC 8739 as reference strain and Saramis^™^ 3.5 (bioMérieux) for spectra interpretation.

### Strain typing

Antibiotic susceptibility of all *E. coli* isolates was tested by broth microdilution using Sensititre COMPAN1F plates (Thermo Fisher Scientific, Waltham, MA, USA) and interpretation according to the Clinical and Laboratory Standards Institute (CLSI) guidelines (14). *E. coli* isolates from pets were then compared to the UTI-causing strains by pulsed-field gel electrophoresis (PFGE) analysis after digestion of total chromosomal DNA with XbaI (New England BioLabs, Ipswich, MA, USA) (15). *Salmonella enterica* serovar Braenderup H9812 was included as an internal control in all gels, and run conditions were as previously described (15). Band patterns were visually compared to define indistinguishable and closely related subtypes within pet/patient pairs according to the criteria proposed by Tenover et al. (16).

In case both a patient and his pet had a 3GC-resistant *E. coli*, both isolates were further characterized by PCR and sequencing for presence of the most common ESBL genes (ɓla_TE_M, ɓla_SHV_, ɓla_CTX-M_) and the plasmid-mediated AmpC gene ɓla_CMY-2_ using previously described primers and protocols (17, 18). Plasmids harboring ESBL or AmpC genes were extracted and transformed into Genehog *E. coli*, followed by PCR-based replicon typing (PBRT) using a commercial kit (Diatheva, Cartoceto, Italy), and S1 nuclease PFGE for size determination, according to previously described protocols (19).

### Follow-up study on patients and pets sharing indistinguishable strains

Patients sharing an *E. coli* strain with their pet based on PFGE were invited to participate in a follow-up study approximately 10 months after their UTI incident. The patients were instructed to collect fecal samples from themselves and their pet. Samples were shipped in plastic containers and arrived at the laboratory the day after collection. On the arrival day, the samples were quantitatively assessed for presence of the original UTI-causing strain by plating 100 μl of serial 10-fold dilutions on MacConkey agar supplemented with the antibiotics corresponding to the resistance profile of the strain. Total coliform counts were assessed by plating the serial dilutions on plain MacConkey agar. Upon overnight incubation, a weighted average count of total and resistant coliform (lactose-positive) colonies was calculated. When present, six *E. coli* colonies from antibiotic-containing plates were subcultured and stored at −80°C.

Human and pet isolates growing on the same antibiotic-containing plates were subjected to whole-genome sequencing. DNA was extracted from the isolates using MasterPure Gram Positive DNA Purification Kit (Epicentre, USA). DNA libraries were prepared using the Nextera XT kit (Illumina Inc., San Diego, CA, USA) and sequenced on the V3 (2 × 300bp) flow cell on the Illumina MiSeq platform, to produce paired-end reads. Genomes were assembled using SPAdes (20) and annotated using RAST (21). Core genes for each set of related isolates were identified using GET_HOMOLOGUES (22). Core gene sequences were aligned using MUSCLE (23) and concatenated to produce an aligned coregenome. The aligned core-genome sequences were the input to produce a maximum-likelihood tree via RAxML (24), with 100 bootstrapping replicates. The number of single nucleotide polymorphisms (SNPs) between isolate pairs was determined from the aligned core-genome. All genomes were screened for their multi-locus sequence (MLST) type and for presence of virulence genes (25), antibiotic resistance genes (26), and plasmid replicons (27) by using the online tools available at the Center for Genomic Epidemiology (https://cge.cbs.dtu.dk/services/).

### Accession numbers

The assembled genomes of seven representative *E. coli* isolates (one per sample) have been deposited in GenBank under accession numbers PXWC00000000 (patient of pair A, UTI-causing isolate), PXVY00000000 (patient of pair A, follow-up fecal sample), PXVZ00000000 (dog of pair A, first fecal sample), PXVX00000000 (dog of pair A, follow-up fecal sample), PXWB00000000 (patient of pair B, UTI-causing isolate), PXWA00000000 (dog of pair B, first fecal sample), and PXVW00000000 (dog of pair B, follow-up fecal sample).

## RESULTS

Among the 119 eligible patients we contacted, nineteen (16%) were pet owners and agreed to participate in the study. Seven of the patients were infected with a 3GC-resistant *E. coli*. Two of the patients lived with two pets, leading to a total of 21 pets sampled, including six cats and 15 dogs.

Antimicrobial selective culture indicated potential strain sharing with a household pet for six of the 12 patients infected with a 3GC-susceptible *E. coli*, and for one of the seven patients infected with a 3GC-resistant strain. For two of these seven patient/pet pairs, the UTI-causing isolate had the exact same antibiotic susceptibility pattern as the corresponding pet isolate (Pairs A and B, Table 1). The isolates within these two patient/pet pairs were also indistinguishable genetically as evidenced by PFGE analysis (Table 1). Pair A comprised a 69-year old woman and her dog that had lived together for three years. Based on data collected through a questionnaire (data not shown), the woman reported daily kissing/licking by the dog, no sharing of furniture, and less than weekly sharing of food. The other pair (pair B) comprised a 53-year old woman and her dog that had been living together for nine years. The woman reported less than weekly kissing/licking by the dog, daily feeding of table scraps, and that the dog was allowed in furniture such as bed and sofa. Canine and human isolates from the single patient/dog pair sharing a 3GC-resistant strain (Pair C) harbored the same ESBL gene (ɓla_CTX-M-15_). However, the canine fecal isolates displayed a different PFGE type compared to the UTI-causing strain, and harbored ɓla_CTX-M-15_ on a 104kb plasmid that was non-typeable by PBRT, whereas the UTI-causing strain carried ɓla_CTX-M-15_ on a 138 kb plasmid that was positive for replicon types FIA, FIB and FII.

**Table 1.** Characterization of *E. coli* isolates from the seven patient/pet pairs for which antimicrobial selective culture indicated potential strain sharing.

| Patient/pet<br>pair | Host<br>(age/gender)<sup>a</sup> | Characterization of *E. coli*<sup>c</sup> |  |  |
| --- | --- | --- | --- | --- |
|  |  | Antimicrobial resistance<sup>b</sup> | ESBL<br>phenotype | PFGE Type |
| A | H (69/♀) | AMP | - | 1 |
|  | D (2/♂) | AMP | - | 1 |
| B | H (53/♀) | AMP, SXT | - | 2 |
|  | D (8/♀) | AMP, SXT | - | 2 |
| C | H (66/♀) | (AMC), AMP, CFZ, CPD, (DOX), MAR, SXT | + | 3 |
|  | D (12/♂) | AMP, CFZ, CPD, MAR, SXT | + | 4 |
| D | H (74/♀) | AMP, SXT | - | 5 |
|  | D (11/♂) | None | - | 6 |
| E | H (74/♀) | (FOX), DOX, SXT | - | 7 |
|  | C (5/♂) | None | - | 8 |
| F | H (67/♀) | (DOX), SXT | - | 9 |
|  | D (1/♂) | SXT | - | 10 |
| G | H (44/♀) | (AMC), AMP, (DOX), GEN, SXT | - | 11 |
|  | D (4/♀) | None | - | 12 |

<sup>a</sup>H, human; D, dog; C, cat. Age is displayed in years.

^a^H, human; D, dog; C, cat. Age is displayed in years.

^b^AMC, amoxicillin/clavulanic acid; AMP, ampicillin; CPD, cefpodoxime; CFZ, cefazolin; DOX, doxycycline; GEN, gentamicin; MAR, marbofloxacin; SXT, trimethoprim/sulfamethoxazole. Brackets indicate intermediate susceptibility.

^c^The human *E. coli* are clinical isolates from UTI’s. The pet *E. coli* are commensal isolates from fecal samples that were collected within 2 weeks after their owner’s UTI episode.

In the follow-up study, the patients of pairs A and B submitted fecal samples from themselves and their dogs approximately 10 months after their UTI incident. Based on susceptibility profiles of the two UTI-causing strains, counts of resistant coliforms were performed on MacConkey agar supplemented with ampicillin (8 μg/ml) in combination with trimethoprim (32 μg/ml) for pair A, and in combination with sulfadiazine (256 μg/ml) for pair B. Coliforms displaying the ampicillin/trimethoprim resistance profile of the UTI-causing strain in pair A were more abundant in the feces of the patient (2.6*10^8^ CFU/g, 100% of total coliforms) than in the feces of her dog (4.3*10^4^ CFU/g, 0.01% of total coliforms). In pair B, coliforms displaying the ampicillin/sulfadiazine resistance profile of the UTI-causing strain were only detected in the feces of the dog (4.4*10^5^ CFU/g, 4.6% of total coliforms). Surprisingly, the feces from the patient of pair B did not result in any coliform growth on plain MacConkey agar.

Whole-genome sequence analysis was performed on 14 isolates from pair A, including the UTI-causing strain, six fecal isolates from the patient and seven fecal isolates from the dog, and on eight isolates from pair B, including the UTI-causing strain and 7 fecal isolates from the dog. The identified SNPs form the basis of the phylogenetic tree in Fig. 1. There was a maximum of 69 and 58 SNPs between any two isolates from pairs A and B, respectively. In pair A, the UTI-causing strain differed from the initial dog fecal isolate by only 5 SNPs, whereas it differed from the follow-up fecal isolates from the patient and dog by 4-13 SNPs and 61-67 SNPs, respectively. Additionally, two integrated phages were identified in all canine follow-up isolates from pair A (Fig. 1). In pair B, the UTI-causing strain differed from the initial dog isolate by 20 SNPs, whereas it differed from the canine follow-up fecal isolates by 22-58 SNPs. All human and canine isolates from pair A belonged to sequence type (ST)80 and harbored *aadAl*, bla_TEM-1A_ and *dfrÄ1* on a Tn7 transposon located on the chromosome (Fig. 2). These isolates also contained the 10 virulence genes *cnf1, gad, iss, iroN, mchB, mchC, mchF, mcmA, pic*, and *vat* (Table 2). All isolates from pair B belonged to ST998 and harbored bla_TEM-1B_, *strAB* and *sul2* on a 11-kb colE1-like plasmid (pPD) (Fig. 2). Additionally, pair B isolates harbored the six virulence genes *cnf1, gad, iroN, iss, sfaS*, and *vat* (Table 2).

**Figure 1.**
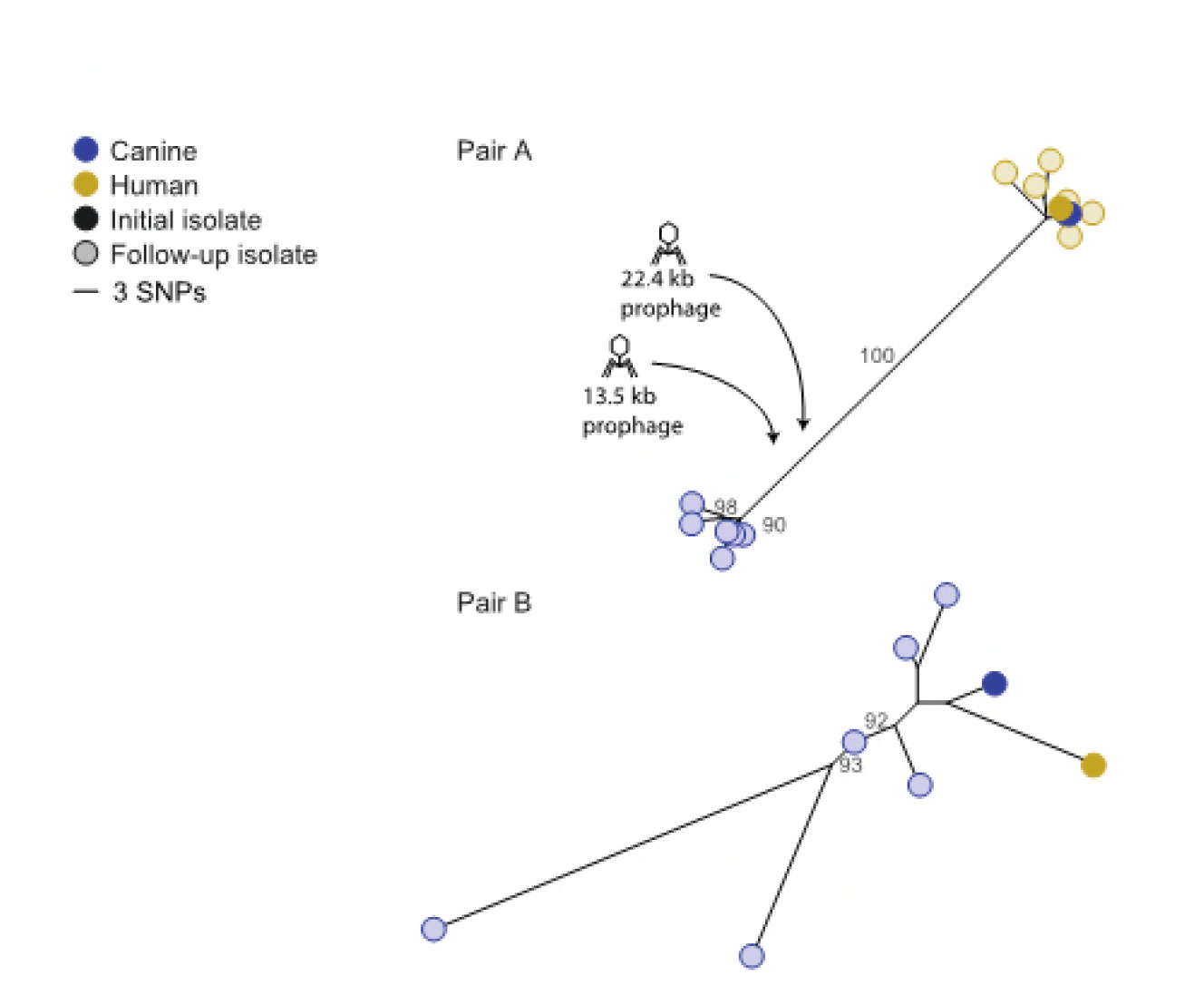
Phylogenetic trees based on SNP data for dog-owner pairs A and B. Solid circles represent isolates taken at the time of the urinary tract infection, whereas lightly-colored circles represent follow-up isolates taken approximately 10 months later. Bootstrap percentages are indicated by the grey number on branches when above 80%. Two integrated phages were identified in the follow-up canine isolates from pair A of indicated sizes.

**Figure 2.**
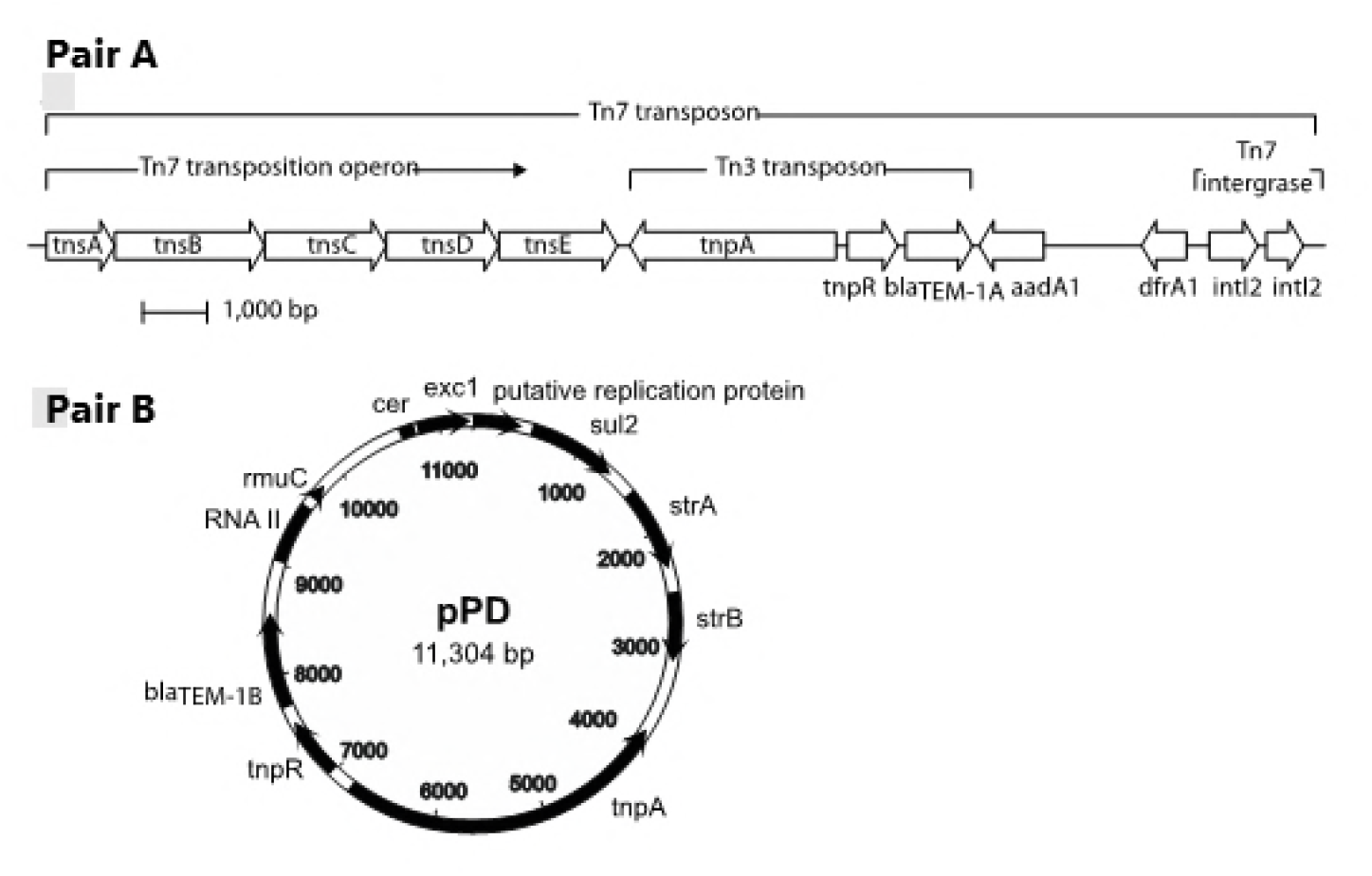
Genetic contexts of all antibiotic resistance genes identified. In isolates of pair A, the resistance genes are located within a Tn7 transposon located on the chromosome. In isolates of pair B, the resistance genes are located on a 11,3 kbp colE1-like plasmid termed pPD.

**Table 2.** Origin and analysis of 22 genome-sequenced isolates obtained from patient/pet pairs A and B at the time of the patients’ UTI infection and approximately 10 months later.

| Patient | Host | Number of | Month/ | Clinical UTI (U) | Multilocus | Virulence genes | Resistance |
| --- | --- | --- | --- | --- | --- | --- | --- |
| /pet |  | isolates | year of | or fecal | sequence |  | genes |
| pair |  | sequenced | isolation | commensal (F) | type |  |  |
|  |  |  |  | origin |  |  |  |
| A | Patient | 1 | 03/2014 | U | ST80 | *cnf1, gad, iroN, iss, mchB, mchC, mchF, mcmA, pic, vat* | *aadA1, bla<sub>TEM-1A</sub>, dfaA1* |
|  |  | 6 | 01/2015 | F |  |  |  |
|  | Dog | 1 | 03/2014 | F |  |  |  |
|  |  | 6 | 01/2015 | F |  |  |  |
| B | Patient | 1 | 03/2014 | U | ST998 | *cnf1, gad, iroN, iss, vat, sfaS,* | *bla<sub>TEM-1B</sub>, strAB, sul2* |
|  | Dog | 1 | 03/2014 | F |  |  |  |
|  |  | 6 | 02/2015 | F |  |  |  |

## DISCUSSION

To our knowledge, this is the first longitudinal study investigating the role of household pets as reservoirs of *E. coli* strains causing UTI in their owners using a quantitative approach. In approximately 10% of the UTI patients, the strain causing infection was detected in the feces of their dog. The two dog-owner pairs that shared the same strain were re-sampled approximately 10 months after the UTI episode, providing insights into strain carriage and possible transmission. For pair A, the relatively higher counts of coliforms displaying the resistance profile of the UTI-causing strain in the patient’s feces than in the dog’s feces suggest that the patient was likely self-infected with a strain that was dominant in her feces, as it is usually the case for UTI patients (2). On the contrary, the UTI-causing strain in pair B was detected only in the dog 10 months after the UTI incident, indicating that this dog was persistent carrier of the strain and a likely source of infection. In that regard, it should be noted that we failed to detect any coliforms in the fecal sample obtained from the patient of pair B. Although the follow-up fecal sample was allegedly processed the day after sampling, we cannot exclude an unreported delay in shipping or improper handling of the sample that may have resulted in bacterial death prior to culture. Alternatively, absence of coliforms in the patient’s feces could be due to a recent antibiotic treatment, but this information was not available. According to the information we received on dog-human interaction, transmission was possible through kissing/licking in pair A and via contamination of the bed and sofa in pair B. In both cases, acquisition of the strain from a common food source cannot be ruled out, since the dogs were periodically fed with food consumed by the owners.

The limited genomic differences observed between human and canine isolates from these two pairs confirmed that the strain residing in the intestinal tract of the dog was the same clone causing UTI in the owner. In pair A, the UTI-causing strain, the first fecal isolate from the dog, and the follow-up fecal isolates from the patient clustered separately from the follow-up fecal isolates from the dog (Fig. 1). This suggests a possible adaptive micro-evolution of the strain in the intestinal tract of the dog during the 10 months elapsing between the sampling times. Such evolution included acquisition of two prophages that could not be detected in the isolates from the owner. Further research is warranted to determine whether these phages may enhance fitness of *E. coli* in the canine intestinal tract. A certain degree of genomic micro-evolution (up to 46 SNPs) also occurred between the first isolate and the follow-up isolates from the dog in pair B (Fig. 1), even though no phage insertions were observed in this case.

The identified resistance genes correlated well with the observed phenotypes (Tables 1 and 2). All isolates from pair A had genes conferring resistance to ampicillin (ɓla_TEM-1A_) and trimethoprim (*dfrAl*), and displayed resistance to these drugs. All isolates from pair B had genes conferring resistance to ampicillin (ɓla_TEM-1B_) and sulfadiazine (*sul2*), and displayed resistance to these drugs. As expected, we detected various ExPEC-associated virulence factors, including an adhesin (*sfaS*), a siderophore receptor involved in iron scavenging (*iroN*), a toxin (*cnf-1*), and a protectin involved in serum resistance (iss) (3, 5). However, the two UTI-causing strains did not contain any of the virulence markers that have been proposed for ExPEC classification (*papA, papC, sfa/foc, afa/dra, iutA* and *kpsMT*) (28). This finding underlines the complexity of defining the ExPEC pathotype, which has a considerable overlap with commensal *E. coli* strains, and overall is much more heterogeneous than other *E. coli* pathotypes, such as those involved in intestinal disease (29).

Among the seven pets screened selectively for 3GC-resistant *E. coli*, one dog shed CTX-M-15-producing *E. coli* as the UTI-causing strain (pair C). However, the canine and the human isolates were genetically unrelated, and the bla_CTX-M-15_ gene was located on different plasmids. This suggests that the dog was not directly implicated in the UTI of the owner but does not exclude that bla_CTX-M-15_ might have transferred between the two hosts prior to insertion into different plasmids. In that regard, frequent insertion of bla_CTX-M-15_ linked to ISEcp1 has been hypothesized based on sequence analysis of CTX-M-15-plasmids in clinical *E. coli* of human and animal origin (30). CTX-M-15 accounts for the majority of the ESBLs found among *E. coli* isolates from infections such as UTI (31). This also occurs as one of the most frequent ESBL-types in *E. coli* isolates from dogs (32), and a recent study has documented sharing of CTX-M-15-producing *E. coli* ST131 by a pet and a child living in the same family household (11).

In conclusion, we have shown definite co-carriership of UTI-causing *E. coli* strains between humans and their pets, and the data indicated one dog as a persistent carrier and a likely source of human infection. Nevertheless, we have not provided a final proof of transmission between the two hosts. Even comprehensive longitudinal studies, where pets and their owners are sampled weekly or monthly, would only provide indications without excluding other possible sources of transmission such as food. Proving a direct pet-associated risk for UTI in owners may be almost impossible, since a number of other risk factors must be at play for UTI to develop. Irrespectively, considering the data presented here and existing knowledge about sharing of bacterial strains between humans and pets, a note of caution could be issued to pet owners not to fraternize too closely with their pets. This would be particularly relevant for pet owners that for some reason have a compromised immune system.

## ACKNOWLEDGMENTS

We greatly acknowledge the technical staff at Hvidovre Hospital for assisting with the identification of eligible patient candidates. This research received no specific grant from any funding agency in the public, commercial, or not-for-profit sectors.

## REFERENCES

1. Ronald A. 2003. The etiology of urinary tract infection: traditional and emerging pathogens. Dis Mon 49:71-82.

2. Yamamoto S, Tsukamoto T, Terai A, Kurazono H, Takeda Y, Yoshida O. 1997. Genetic evidence supporting the fecal-perineal-urethral hypothesis in cystitis caused by *Escherichia coli*. J Urol 157:1127-1129.

3. Johnson J, Russo TA. 2002. Extraintestinal pathogenic *Escherichia coli*: “the other bad *E coli”*. J Lab Clin Med 139:155-162.

4. Russo TA Johnson JR. 2000. Proposal for a new inclusive designation for extraintestinal pathogenic isolates of *Escherichia coli*: ExPEC. J Infect Dis 181:1753-1754.

5. Johnson JR, Stell AL, Delavari P. 2001. Canine feces as a reservoir of extraintestinal pathogenic *Escherichia coli*. Infect Immun 69:1306-1314.

6. Low DA, Braaten BA, Ling GV, Johnson DL, Ruby AL. 1988. Isolation and comparison of *Escherichia coli* strains from canine and human patients with urinary tract infections. Infect Immun 56:2601-2609.

7. Platell JL, Cobbold RN, Johnson JR, Clabots CR, Trott DJ. 2012. Fluoroquinolone-resistant extraintestinal *Escherichia coli* clinical isolates representing the O15:K52:H1 clonal group from humans and dogs in Australia. Comp Immunol Microbiol Infect Dis 35:319-324.

8. Damborg P, Nielsen SS, Guardabassi L. 2009. *Escherichia coli* shedding patterns in humans and dogs: insights into within-household transmission of phylotypes associated with urinary tract infections. Epidemiol Infect 137:1457-1464.

9. Johnson JR, Owens K, Gajewski A, Clabots C. 2008. *Escherichia coli* colonization patterns among human household members and pets, with attention to acute urinary tract infection. J Infect Dis 197:218-224.

10. Johnson JR, Clabots C. 2006. Sharing of virulent *Escherichia coli* clones among household members of a woman with acute cystitis. Clin Infect Dis 43:e101-108.

11. Johnson JR, Davis G, Clabots C, Johnston BD, Porter S, DebRoy C, Pomputius W, Ender PT, Cooperstock M, Slater BS, Banerjee R, Miller S, Kisiela D, Sokurenko EV, Aziz M, Price LB. 2016. Household clustering of *Escherichia coli* sequence type 131 clinical and fecal isolates according to whole genome sequence analysis. Open Forum Infect Dis 3:ofw129.

12. Murray AC, Kuskowski MA, Johnson JR. 2004. Virulence factors predict *Escherichia coli* colonization patterns among human and animal household members. Ann Intern Med 140:848-849.

13. Bartoloni A, Bartalesi F, Mantella A, Dell’Amico E, Roselli M, Strohmeyer M, Barahona HG, Barrón VP, Paradisi F, Rossolini GM. 2004. High prevalence of acquired antimicrobial resistance unrelated to heavy antimicrobial consumption. J Infect Dis 189:1291-1294.

14. Clinical Laboratory Standards Institute (CLSI). Performance standards for antimicrobial disk and dilution susceptibility tests for bacteria isolated from animals; second informational supplement. CLSI document VET01S2. Wayne: CLSI; 2013.

15. Ribot EM, Fair MA, Gautom R, Cameron DN, Hunter SB, Swaminathan B, Barrett TJ. 2006. Standardization of pulsed-field gel electrophoresis protocols for the subtyping of *Escherichia coli* O157:H7, *Salmonella*, and *Shigella* for PulseNet. Foodborne Pathog Dis 3:59-67.

16. Tenover FC, Arbeit RD, Goering RV, Mickelsen PA, Murray BE, Persing DH, Swaminathan B. 1995. Interpreting chromosomal DNA restriction patterns produced by pulsed-field gel electrophoresis: criteria for bacterial strain typing. J Clin Microbiol 33:2233-2239.

17. Hasman H, Mevius D, Veldman K, Olesen I, Aarestrup FM. 2005. Beta-lactamases among extended-spectrum beta-lactamase (ESBL)-resistant *Salmonella* from poultry, poultry products and human patients in The Netherlands. J Antimicrob Chemother 56:115-121.

18. Pérez-Pérez FJ, Hanson ND. 2002. Detection of plasmid-mediated AmpC beta-lactamase genes in clinical isolates by using multiplex PCR. J Clin Microbiol 40:2153-2162.

19. Hansen KH, Bortolaia V, Damborg P, Guardabassi L. 2014. Strain diversity of CTX-M-producing Enterobacteriaceae in individual pigs: insights into the dynamics of shedding during the production cycle. Appl Environ Microbiol 80:6620-6626.

20. Bankevich A, Nurk S, Antipov D, Gurevich AA, Dvorkin M, Kulikov AS, Lesin VM, Nikolenko SI, Pham S, Prjibelski AD, Pyshkin AV, Sirotkin AV, Vyahhi N, Tesler G, Alekseyev MA, Pevzner PA. 2012. SPAdes: a new genome assembly algorithm and its applications to single-cell sequencing. J Comput Biol 19:455-477.

21. Aziz RK, Bartels D, Best AA, DeJongh M, Disz T, Edwards RA, Formsma K, Gerdes S, Glass EM, Kubal M, Meyer F, Olsen GJ, Olson R, Osterman AL, Overbeek RA, McNeil LK, Paarmann D, Paczian T, Parrello B, Pusch GD, Reich C, Stevens R, Vassieva O, Vonstein V, Wilke A, Zagnitko O. 2008. The RAST Server: rapid annotations using subsystems technology. BMC Genomics 9:75.

22. Contreras-Moreira B, Vinuesa P. 2013. GET_HOMOLOGUES, a versatile software package for scalable and robust microbial pangenome analysis. Appl Environ Microbiol 79:7696-7701.

23. Edgar RC. 2004. MUSCLE: multiple sequence alignment with high accuracy and high throughput. Nucleic Acids Res 32:1792-1797.

24. Stamatakis A. 2014. RAxML version 8: a tool for phylogenetic analysis and post-analysis of large phylogenies. Bioinformatics 30:1312-1313.

25. Joensen KG, Scheutz F, Lund O, Hasman H, Kaas RS, Nielsen EM, Aarestrup FM. 2014. Real-time whole-genome sequencing for routine typing, surveillance, and outbreak detection of verotoxigenic *Escherichia coli*. J Clin Microbiol 52:1501-1510.

26. Zankari E, Hasman H, Cosentino S, Vestergaard M, Rasmussen S, Lund O, Aarestrup FM, Larsen MV. 2012. Identification of acquired antimicrobial resistance genes. J Antimicrob Chemother 67:2640-2644.

27. Carattoli A, Zankari E, García-Fernández A, Voldby Larsen M, Lund O, Villa L, Møller Aarestrup F, Hasman H. 2014. In silico detection and typing of plasmids using PlasmidFinder and plasmid multilocus sequence typing. Antimicrob Agents Chemother 58:3895-3903.

28. Johnson JR, Murray AC, Gajewski A, Sullivan M, Snippes P, Kuskowski MA, Smith KE. 2003. Isolation and molecular characterization of nalidixic acid-resistant extraintestinal pathogenic *Escherichia coli* from retail chicken products. Antimicrob Agents Chemother 47:2161-2168.

29. Köhler CD, Dobrindt U. 2011. What defines extraintestinal pathogenic *Escherichia coli*? Int J Med Microbiol. 301:642-647.

30. Smet A, Van Nieuwerburgh F, Vandekerckhove TT, Martel A, Deforce D, Butaye P, Haesebrouck F. 2010. Complete nucleotide sequence of CTX-M-15-plasmids from clinical *Escherichia coli* isolates: insertional events of transposons and insertion sequences. PLoS One 5:e11202.

31. Cantón R, González-Alba JM, Galán JC. 2012. CTX-M enzymes: origin and diffusion. Front Microbiol 3:110.

32. Ewers C, Bethe A, Semmler T, Guenther S, Wieler LH. 2012. Extended-spectrum β-lactamase-producing and AmpC-producing *Escherichia coli* from livestock and companion animals, and their putative impact on public health: a global perspective. Clin Microbiol Infect 18:646-655.

